# Haplotype Explorer: an infection cluster visualization tool for spatiotemporal dissection of the COVID-19 pandemic

**DOI:** 10.1101/2020.07.19.179101

**Authors:** Tetsuro Kawano-Sugaya, Koji Yatsu, Tsuyoshi Sekizuka, Kentaro Itokawa, Masanori Hashino, Rina Tanaka, Makoto Kuroda

**Affiliations:** Pathogen Genomics Center, National Institute of Infectious Diseases, Tokyo, Japan

**Keywords:** SARS-CoV-2, COVID-19, Haplotype network, Infection clusters, Infectious diseases, epidemiology

## Abstract

The worldwide eruption of COVID-19 that began in Wuhan, China in late 2019 reached 10 million cases by late June 2020. In order to understand the epidemiological landscape of the COVID-19 pandemic, many studies have attempted to elucidate phylogenetic relationships between collected viral genome sequences using haplotype networks. However, currently available applications for network visualization are not suited to understand the COVID-19 epidemic spatiotemporally, due to functional limitations That motivated us to develop Haplotype Explorer, an intuitive tool for visualizing and exploring haplotype networks. Haplotype Explorer enables people to dissect epidemiological consequences via interactive node filters to provide spatiotemporal perspectives on multimodal spectra of infectious diseases, including introduction, outbreak, expansion, and containment, for given regions and time spans. Here, we demonstrate the effectiveness of Haplotype Explorer by showing an example of its visualization and features. The demo using SARS-CoV-2 genome sequences is available at https://github.com/TKSjp/HaplotypeExplorer

**Summary:** A lot of software for network visualization are available, but existing software have not been optimized to infection cluster visualization against the current worldwide invasion of COVID-19 started since 2019. To reach the spatiotemporal understanding of its epidemics, we developed Haplotype Explorer. It is superior to other applications in the point of generating HTML distribution files with metadata searches which interactively reflects GISAID IDs, locations, and collection dates. Here, we introduce the features and products of Haplotype Explorer, demonstrating the time-dependent snapshots of haplotype networks inferred from total of 4,282 SARS-CoV-2 genomes.

## Introduction

To eliminate infectious diseases, it is essential to quickly identify and control emerging infection clusters before they become uncontrollable. For this purpose, many applications to assist researchers in understanding the latest epidemiology have been developed. In fact, the recent intensification of the COVID-19 pandemic, which began in late 2019 in Wuhan, China, has prompted development of new software to support investigations of this virus. For example, Nextstrain (Hadfield et al. 2018) is one of the most popular web services related to the COVID-19 pandemic, and provides interactive molecular phylogenetic trees and geographic maps representing possible virus transmission routes. COVID-19 Genome Tracker (Akther et al. 2020) is another unique application which shows the evolution of SARS-CoV-2 using a haplotype network. This tool can dynamically display metadata, such as isolate conditions, locations, and mutations, compared to the reference genome.

So far, many phylogenetic trees and haplotype networks using the SARS-CoV-2 genome have been inferred using Nextstrain and COVID-19 Genome Tracker because they are suited to interpret genetic and epidemiological relationships among sequences. Haplotype networks are especially useful for this pandemic due to their potential for displaying short-term diversification of closely-related genomes. Many available software programs for network inferring, such as TCS (Clement et al. 2000), PopART (Leigh and Bryant 2015), and Network (Bandelt et al. 1999), have supported these studies using haplotype networks of SARS-CoV-2. Although these applications also work as network viewers, many alternatives are also available for additional annotation and exploration, including Cytoscape (Su et al. 2014), Gephi Bastian et al. 2009), and tcsBU (Múrias dos Santos et al. 2016).

Despite the excellent designs of the above-mentioned tools, none of the currently available software programs simultaneously fulfill the requirements essential to dissect epidemic situations: 1) nodes that can be dynamically filtered with metadata by complex search queries, 2) nodes that can be indicated by real-time pie charts which reflect sample size and content proportions at a given point in time, and 3) interactive distributions requiring no external software installation other than a modern web browser. Hence, we endeavored to develop Haplotype Explorer, a specialized network viewer, to assist onsite actions against emerging pathogens.

## Methods

### Implementation and workflow

Haplotype Explorer is a JavaScript application executable in web browser, so it does not require uploading data to an external web server. This allows users to analyze confidential data securely. It can produce distributions in HTML, which enables users to share originated networks with others easily. The network structure is written in JavaScript Object Notation (JSON) format, and can be generated semi-automatically from a multi-FASTA file with the provided programs. The flow-diagram starting from the raw multi-FASTA file to resultant network graphs is shown in Figure 2. Briefly, the procedure is as follows: 1) collect, curate, and align sequences using the provided python programs and other external programs, 2) run network analysis in TCS, and 3) convert data to an HTML file. All procedures are supported by the provided python programs (Step1-3.py). We confirmed compatibilities of Haplotype Explorer and the bundled python scripts with the latest versions of Safari, Firefox, Edge, Chrome, and Python3 on macOS Catalina 10.15.3, respectively.

### Data analysis

Whole genome sequences were retrieved from GISAID (Shu and McCauley 2017) on 9 June 2020 using the following options: 1) collection date was before 21 March 2020, 2) host was only human, 3) check was on for “complete,” “high coverage,” and “low coverage excl.” After retrieval of a total of 9,583 sequences, they were curated by Step1.py. Briefly, the Step1.py is in charge of removing low-quality sequences, such as those containing spaces, gaps, degenerated bases, and ambiguous collection dates (i.e., month or date are absent in collection date) using seqkit (Shen et al. 2016) and the Linux sed command. Passing sequences were aligned by MAFFT Katoh et al. 2002), clustered by CD-HIT (Fu et al. 2012) (threshold: 100% identical), and SNVs extracted by snp-sites Page et al. 2016). The Step1.py output PHYLIP file is an input of TCS, and the Step2.py assists in executing TCS. The resultant GraphML (.gml) file was parsed by the Step3.py to generate the resultant HTML file for network exploration. In the following analysis, we collected figures by applying the filters “~YYYYMMDD” day by day from the initial day (31 December 2019; Wuhan-Hu-1) to 21 March 2020.

## Results and Discussion

### Principal features of Haplotype Explorer toward epidemic dissection

The primary feature of Haplotype Explorer is a vibrant and interactive visualization function utilizing D3.js (Bostock et al. 2011) and metadata, including sample size, accession no, location, and collection date, which are important to understand epidemics. Each node is represented by differently-sized pie charts calculated from sample size and location proportion described in the metadata. The node size is automatically calculated using metadata. The nodes and related edges can be interactively highlighted when a specific node is left-clicked, making it easy to dissect a crowded network with large samples (Figure 1). Users can quickly look into the node of interest by zooming with the scroll-wheel, and show metadata by mousing-over the tool-tip window. The application has four text boxes for filtering nodes: GISAID ID, location, YYYYMMDD~, and ~YYYYMMDD. Filters can be combined, and the GISAID ID and location can be specified by regular expressions. The current view of the network can be exported in a JSON format file, and users can resume it by importing the JSON. Finally, the current SVG view can be converted into a high-resolution PNG image using the export button, which is able to be saved by left-clicking. We also provide python scripts for assisting haplotype network construction with in-house data. Details are shown in the flow-diagram (Figure 2).

**Figure 1.**
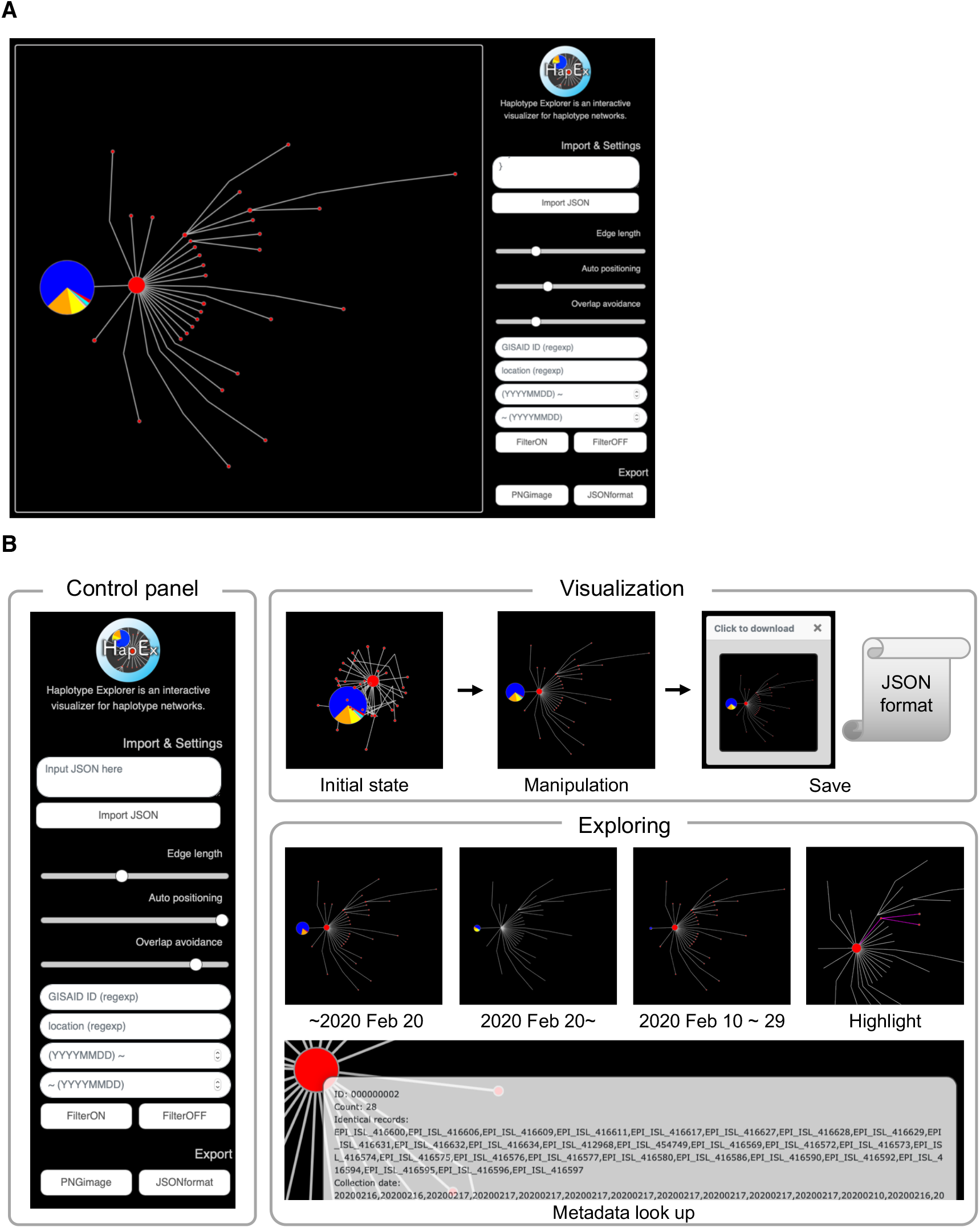
Introduction of Haplotype Explorer. **(A)** The view of Haplotype Explorer **(B)** Introduction of features of Haplotype Explorer. It can modify distances and attractive forces among nodes and avoid overlapping automatically using the slide bar. Nodes can be moved by dragging. After manipulation of the appearance of the network, the view can be exported into PNG and JSON formats. Nodes are easily hidden or visible depending on keyword filters; accession ID, location, collection date from YYYYMMDD, and until YYYYMMDD. In cases where users specify dates, the pie chart is redrawn according to metadata so as to match to the queues. The metadata is displayed by mouse-hover, making it easy to inspect the node of interest.

**Figure 2.**
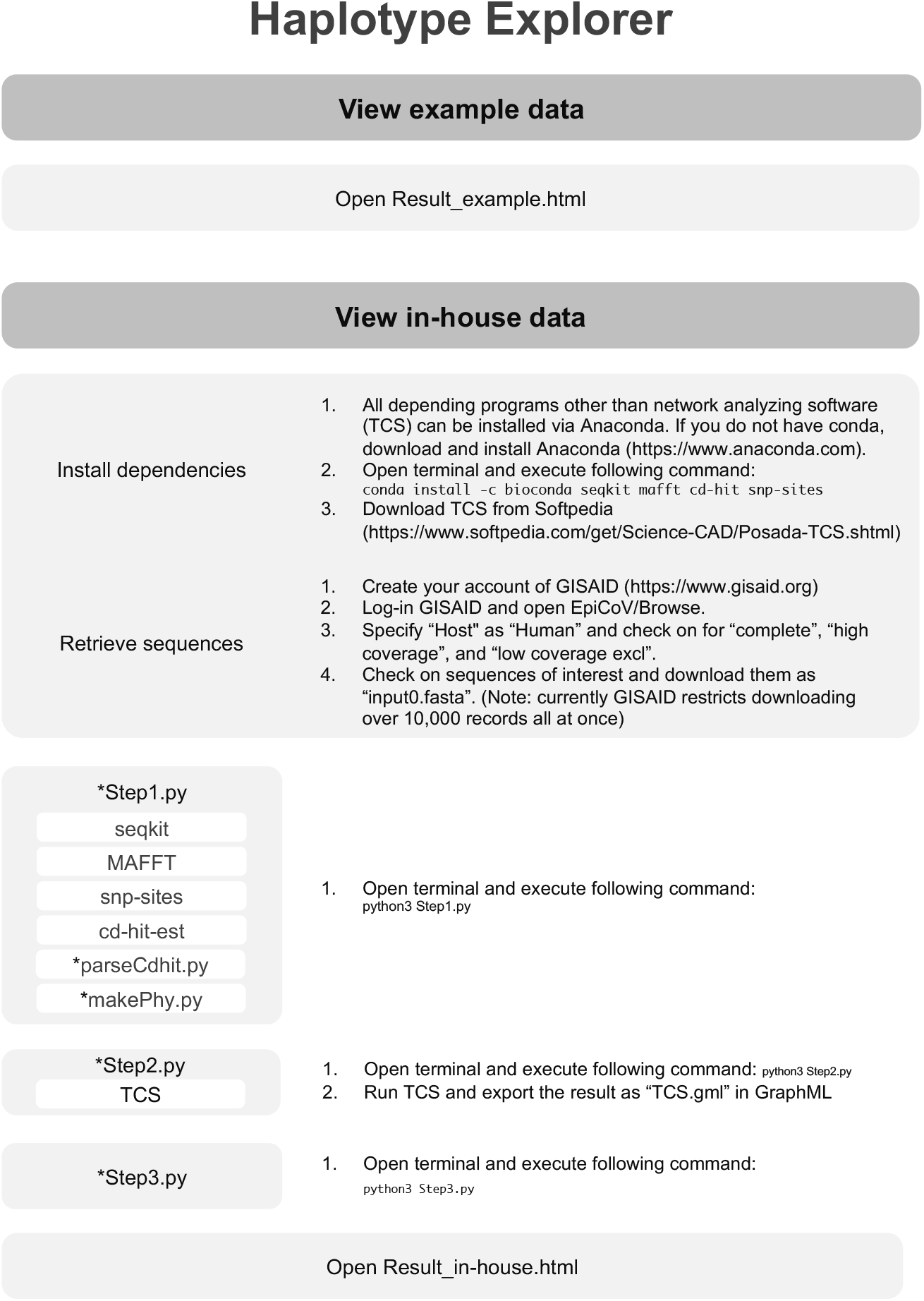
Workflow and dependencies of Haplotype Explorer. The distribution of Haplotype Explorer bundles the example result consisting of the SARS-CoV-2 haplotype network, which can be visualized by opening Result_example.html on a web browser. This viewer can also visualize in-house data using three-step commands. Users can start from Step1.py, which processes in-house multi-fasta (any multi-fasta is available, whereas SARS-CoV-2 genomes retrieved via GISAID are assumed), followed by TCS analysis by Step2.py, and creation of an html file by Step3.py. Asterisks indicate python scripts provided in the current work.

### Demonstration of Haplotype Explorer: spatiotemporal dissection of multimodal epidemics

The epidemic context of SARS-CoV-2 from January 1 to March 21 was visualized by Haplotype Explorer (Figure 3). We began by capturing a snapshot for 21 March 2020 as an overall view. Haplotype Explorer effectively discerned significantly large, but distinctly invaded clusters consisting of a dozen to over one hundred genome collections formed by late March (Figure 3A; magenta arrowhead). In order to understand epidemics in a time-dependent manner, Haplotype Explorer can also generate snapshots for specified dates (Figure 3B).

**Figure 3.**
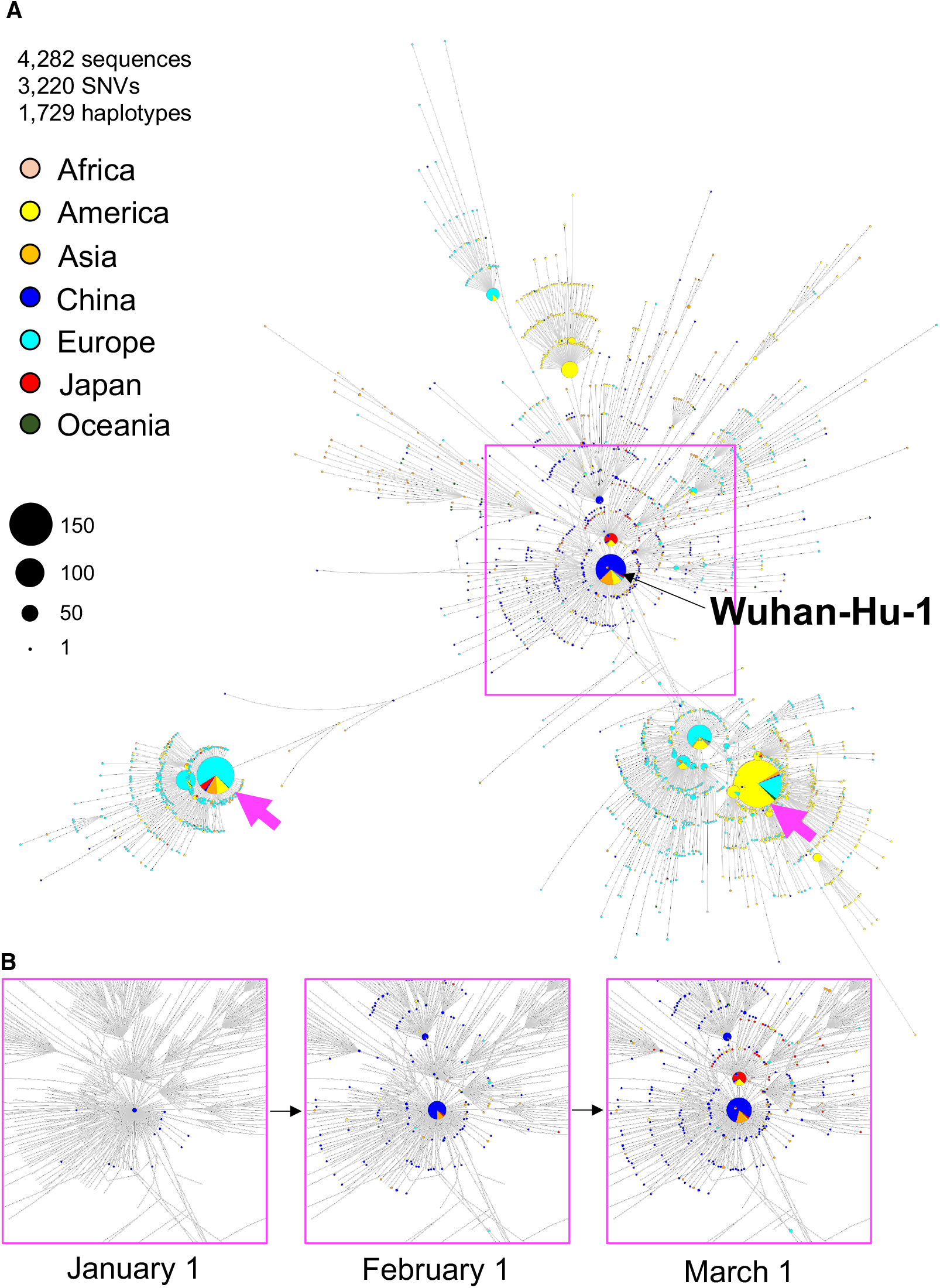
Demonstration of spatiotemporal analysis of the SARS-CoV-2 genomic network using Haplotype Explorer. (A) An example of the exported network generated by Haplotype Explorer using 1,729 of SNVs calculated from 4,282 of world-wide SARS-CoV-2 genomes until 21 March 2020 obtained from the GISAID database. Each node size depends on sample size, and node colors differ by locations. The black arrow indicates the Wuhan-Hu-1 reference sequence. Magenta arrows indicate distinct erosion of SARS-CoV-2 occurring mainly in Europe or America. (B) Three snapshots of the SARS-CoV-2 genomic network around Wuhan, China from January 1 to March 1. Haplotype Explorer enabled us to dissect a haplotype network depending on metadata, giving significant insights into the epidemic.

## Supporting information

Demo of Haplotype Explorer

## Author contributions

Manuscript preparation: TKS MK

Data analysis: TKS MK KY TS

Data collection: KI MH RT

## Funding

This study was supported by a Grant-in Aid from the Japan Agency for Medical Research and Development (AMED) under Grant number JP19fk0108103, JP19fk0108104, and JP20fk0108063. The funding agencies had no role in the study design, data collection or analysis, decision to publish, or manuscript preparation. We would like to thank all authors who have deposited genome sequences in the GISAID EpiCoV Database.

## Conflict of Interest

none.

## Figure legends

**Table.** Comparison of features with other applications. The significance of Haplotype Explorer is that it enables users to generate an explorable HTML distribution, which includes several features all in one. It requires no external software other than a modern web browser to open the network, making it easy to share data. Furthermore, it can draw nodes as pie charts based on the specified span input into the search boxes, supporting spatiotemporal dissection of the network.

|  | Haplotype Explorer | tcsBU<sup>8</sup> | COVID-19 Genome Tracker<sup>2</sup> | Cytoscape<sup>6</sup> | Gephi<sup>7</sup> | PopART<sup>4</sup> | TCS<sup>3</sup> |
| --- | --- | --- | --- | --- | --- | --- | --- |
| Generate explorable HTML distribution | ✓ |  |  |  |  |  |  |
| Time-dependent node pie charts | ✓ |  |  |  |  |  |  |
| Filter by metadata | ✓ |  | ✓ | ✓ | ✓ | ✓ |  |
| Resume operation | ✓ |  |  | ✓ | ✓ | ✓ | ✓ |
| Export image | ✓ | ✓ |  | ✓ | ✓ | ✓ | ✓ |
| Use in-house data | ✓ | ✓ |  | ✓ | ✓ | ✓ | ✓ |
| Real-time orchestration of node positions | ✓ | ✓ | ✓ |  | ✓ |  |  |
| Run on web browser | ✓ | ✓ | ✓ |  |  |  |  |

## Notes

https://github.com/TKSjp/HaplotypeExplorer

### Competing Interest Statement

The authors have declared no competing interest.

## References

1. Hadfield J. et al. (2018) Nextstrain: real-time tracking of pathogen evolution. Bioinformatics. 34(23):4121–4123. doi: https://doi.org/10.1093/bioinformatics/bty407

2. Akther S. et al. (2020) CoV Genome Tracker: tracing genomic footprints of Covid-19 pandemic. bioRxiv. doi: https://doi.org/10.1101/2020.04.10.036343

3. Clement M. et al. (2000) TCS: a computer program to estimate gene genealogies. Mol Ecol. 9(10):1657–1659. doi: https://doi.org/10.1046/j.1365-294x.2000.01020.x

4. Leigh J. W., Bryant D. (2015). PopART: Full-feature software for haplotype network construction. Methods Ecol Evol 6(9):1110–1116

5. Bandelt H. J. et al. (1999) A. Median-joining networks for inferring intraspecific phylogenies. Mol Biol Evol. 16(1):37–48. doi: https://doi.org/10.1093/oxfordjournals.molbev.a026036

6. Su G. et al. (2014). Biological network exploration with Cytoscape 3. Curr Protoc Bioinformatics. 47:8.13.1–8.13.24. doi: https://doi.org/10.1002/0471250953.bi0813s47

7. Bastian M. et al. (2009) Gephi: an open source software for exploring and manipulating networks. International AAAI Conference on Weblogs and Social Media.

8. Múrias dos Santos A. et al. (2016) tcsBU: a tool to extend TCS network layout and visualization. Bioinformatics. 32(4):627–628. doi: https://doi.org/10.1093/bioinformatics/btv636

9. Bostock M. et al. (2011) D^3^Data-Driven Documents. IEEE Transactions on Visualization and Computer Graphics. doi: https://doi.org/10.1109/TVCG.2011.185

10. Shu Y., McCauley J. (2017) GISAID: Global initiative on sharing all influenza data – from vision to reality. Euro Surveill. 22(13) doi: https://doi.org/10.2807/1560-7917.ES.2017.22.13.30494

11. Shen W. et al. (2016) SeqKit: A Cross-Platform and Ultrafast Toolkit for FASTA/Q File Manipulation. PLoS ONE 11(10): e0163962. doi: https://doi.org/10.1371/journal.pone.0163962

12. Katoh K. et al. (2002) MAFFT: a novel method for rapid multiple sequence alignment based on fast Fourier transform. Nucl Acid Res. doi: https://doi.org/10.1093/nar/gkf436

13. Fu L. et al. (2012) CD-HIT: accelerated for clustering the next generation sequencing data. Bioinformatics. 28 (23): 3150–3152. doi: https://doi.org/10.1093/bioinformatics/bts565

14. Page A. J. et al. (2016) SNP-sites: rapid efficient extraction of SNPs from multi-FASTA alignments. Microb Genom. 2(4):e000056. doi: https://doi.org/10.1099/mgen.0.000056

