## Supplementary material for "Haplotype Explorer: an infection cluster visualization tool for spatiotemporal dissection of the COVID-19 pandemic": Demo of Haplotype Explorer: Example-1.html


**Click to download**

×

Haplotype Explorer is an interactive visualizer for haplotype networks.

  

##### Import & Settings

Import JSON
  
  
Edge length
  

Auto positioning
  

Overlap avoidance
  


FilterON
FilterOFF  
  

##### Export

PNGimage
JSONformat
